# Single-cell multi-omics analysis reveals heterogeneity and plasticity of neutrophil states in response to immunotherapies

**DOI:** 10.64898/2026.07.02.735691

**Authors:** Anqi Gao, Sananda Shyamkumar, Natalie B. Winn, Amy K. Erbe, Shelly Davis, Jennifer Zaborek, Kaia Heimstreet, Simon Boyenga, Joel Matthews, Stella Tzu-Ming Tsao, Paul M. Sondel, Huy Q. Dinh

## Abstract

**Background:** Tumor-associated neutrophils (TANs) are emerging as functionally heterogeneous and plastic cells in the tumor microenvironment. In immunologically cold tumors, elevated neutrophil abundance correlates with poor prognosis and resistance to immune checkpoint inhibition (ICI). Whether distinct anti-tumoral neutrophil states can be induced by different immunotherapies and how they relate to treatment efficacy remains unclear.

**Methods:** Using the syngeneic MOC2-huEGFR (M2h) mouse model of head and neck squamous cell cancer (HNSCC), we treated tumor-bearing mice with agonistic anti-CD40 monoclonal antibody (mAb) (aCD40), TNFα, Cetuximab, or a combination of all three, designated Neutrophil Activating Therapy (NAT). In addition to evaluating anti-tumor efficacy, we performed single-cell multiomics RNA and protein sequencing, followed by bioinformatics analyses and flow cytometry validation. NAT-induced anti-tumor efficacy and related neutrophil states were also assessed in another cold tumor model, 9464D-GD2 neuroblastoma. Murine treatment-induced neutrophil gene signatures were then evaluated using clinical, proteomic, and transcriptomic data from HNSCC patients.

**Results:** Five transcriptionally distinct neutrophil states (N0–N4), including precursor state CD49d^+^ N4, were identified using the M2h model. N0 neutrophils (immunosuppressive/quiescent) dominated untreated tumors, but not in successful treatments. ISG^+^ N1 neutrophils and CCR3^+^ N3 neutrophils expanded by aCD40, TNFα, and NAT treatment with anti-tumoral gene signatures and found more interacting with CD8^+^ T cells from bioinformatics analysis. N2 neutrophils reflected a recently established hypoxia-adapted state found in all treatments. ICAM1 (CD54) emerged as a marker of treatment-induced neutrophil activation, discriminating N1, N2, and N3 neutrophils from N0 neutrophils, validated by flow cytometry. In the 9464D-GD2 neuroblastoma model, NAT treatment also reduced the N0 dominance seen in untreated tumors in the HNSCC model but failed to induce anti-tumoral neutrophil states. In 23 HNSCC patients who received ICI therapy, ICAM1 protein expression in neutrophils trended toward association with responder status (TMA-level p=0.029), and ICAM1 neutrophil gene expression also trended toward association with improved overall survival in TCGA data (HR=0.75, p=0.059).

**Conclusions:** Distinct immunotherapy-induced neutrophil states are defined by transcriptional profiles enriched in different functional pathways, associated with both anti-tumor and pro-tumor signatures. ICAM1 identifies activated neutrophils and potentially serves as a biomarker of ICI response in HNSCC, warranting further clinical validation.

**WHAT IS ALREADY KNOWN ON THIS TOPIC:** Neutrophil heterogeneity has received increasing attention, with studies identifying antitumoral neutrophil populations, either at baseline or induced by treatment. Several effective treatment regimens involve an anti-CD40 agonist (aCD40) antibody, among them Neutrophil Activating Therapy (NAT), which combines aCD40, TNFα, and a tumor antigen binding antibody designed to reprogram neutrophils. NAT could thus be particularly effective in cold, myeloid-rich tumors that are largely unresponsive to conventional immunotherapies such as checkpoint blockade, enacting these anti-tumoral effects through similar and different mechanisms; however, this has not been tested.

**WHAT THIS STUDY ADDS:** This study adds a single-cell multi-omics framework for defining treatment-induced neutrophil heterogeneity in MOC2-huEGFR and 9464D-GD2 tumors, two immunologically cold models. It highlights ICAM1/CD54 and interferon-stimulated genes as markers of a dominant antitumor neutrophil state, while showing that neutrophil state composition variy across tumor models.

**HOW THIS STUDY MIGHT AFFECT RESEARCH, PRACTICE, OR POLICY:** These results support the efficacy of a myeloid-modulating therapy built around aCD40 and TNFα in a cold murine head and neck cancer model, and to a lesser extent in a cold murine neuroblastoma model. ICAM1/CD54 expression in neutrophils was also identified as a promising marker of antitumor activity and treatment response. More broadly, this work suggests that incorporating aCD40 and/or TNFα into existing treatment regimens could improve outcomes, while ICAM1/CD54-high neutrophils may serve as a useful therapeutic readout.

## INTRODUCTION

Tumor-associated neutrophils (TANs) are functionally heterogeneous and transcriptionally plastic, challenging earlier views of neutrophils as uniformly pro-tumoral cells in the tumor microenvironment (TME).[1,2] Pan-cancer analyses from The Cancer Genome Atlas (TCGA) RNA-Seq showed that increased neutrophil abundance within tumors is associated with poor prognosis across multiple cancer types.[3] Proposed pro-tumoral mechanisms include suppression of T and NK cell cytotoxicity, promotion of angiogenesis via VEGF and MMP9, and immunosuppressive cytokine production.[4–6] The earlier introduction of N1/N2 nomenclature represented a framework for TAN heterogeneity, distinguishing antitumoral (N-1) from pro-tumoral (N-2) subsets.[6] Advances in single-cell multi-omics profiling technologies have revealed a greater TAN diversity, including emerging pro-tumoral states associated with hypoxia[7], or senescent CCL3^+^ state[8], as well as anti-tumoral interferon-stimulated and antigen-presenting states, each characterized by distinct transcriptional programs and functional implications.[9,10]

Neutrophil plasticity in cancer is governed by spatiotemporal maturation cues, TME-derived signals, including cytokines, hypoxia, and necrotic factors, as well as interactions with tumor cells, other immune populations, and stromal cells.[1] This plasticity is increasingly recognized as both a vulnerability and a therapeutic opportunity. Whereas neutrophils in untreated cold tumors are typically pro-tumoral and immunosuppressive, successful immunotherapy may reprogram them toward anti-tumoral states. Recent work has shown that agonistic anti-CD40 mAb (aCD40) induces interferon-stimulated gene (ISG^+^) anti-tumoral neutrophils in murine lung and colon cancer models[9]; OX40 agonism promotes anti-tumoral TANs in melanoma[11]; and the Neutrophil Activating Therapy (NAT) combining aCD40, TNFα, and a tumor-targeting mAb, drives potent neutrophil reprogramming associated with anti-tumor efficacy across multiple mouse models.[12] Whether these anti-tumoral neutrophil states can be induced in solid tumors that are unresponsive to current immunotherapies, and how treatment-induced neutrophil reprogramming differs across regimens, remains poorly defined.

Here, we studied immunotherapy-induced neutrophil heterogeneity in Head and Neck Squamous Cell Carcinoma (HNSCC), an immunologically cold tumor with a low mutational burden, in which only ∼20% of patients respond to Pembrolizumab, FDA-approved as frontline therapy for recurrent or metastatic disease since 2019.[13] The HNSCC TME is characterized by abundant immunosuppressive myeloid cells, and elevated pretreatment neutrophil-to-lymphocyte ratios (NLR>3–5) consistently correlate with worse progression-free and overall survival among patients receiving ICI.[14,15] Given the limited response to ICI, which primarily leverages T cell functions, there is a need for better and more targeted HNSCC immunotherapies, potentially including approaches that target myeloid cells. Among HNSCC models, the mouse oral cell carcinoma line 2 (MOC2) has been extensively characterized, contains a predominately neutrophilic immune compartment[16], and is non-responsive to ICI[17], making it an attractive model for testing novel immunotherapy combinations. The MOC2-huEGFR (M2h) model recapitulates key immunological features of HNSCC, is nonresponsive to ICI, and expresses human EGFR, enabling evaluation of Cetuximab, an FDA-approved anti-EGFR mAb that binds most HNSCC tumors, as a component of NAT.[18]

In this study, we used M2h to test the hypothesis that myeloid-centric immunotherapy induces distinct neutrophil states with anti-tumoral potential. Using single-cell multi-omics RNA and protein sequencing followed by bioinformatics analysis, we characterize five transcriptionally TAN states (N0–N4) with both pro- and anti-tumoral gene signatures following treatment with NAT. Additionally, cross-model analysis in another immunologically cold tumor model, 9464D-GD2 murine neuroblastoma, further established treatment-associated phenotypic shifts relative to untreated tumors and the context dependence of NAT-induced neutrophil changes across cold tumor types. Using clinical data, we also provide translational evidence that ICAM1 expression, a marker of treatment-induced TANs, is associated with ICI response and improved survival in HNSCC patients.

## METHODS

### Mouse tumor models and treamtment

C57BL/6 female mice (6-8 weeks; Jackson Laboratory or Biotron) were engrafted intradermally with 2×10⁶ M2h cells or 9464D-GD2 cells in 100 μl PBS.[18,19] M2h cells and 9464D-GD2 cells were cultured according to previously published protocols.[18,19] Tumors were measured twice weekly in a blinded manner, and tumor volumes were calculated as (width²×length)/2.[18,19] Randomization of mice to treatment groups was completed at small (35–50 mm³) or large (125–150 mm³) tumor volumes for mice bearing M2h tumors and at small (35-50 mm³) volumes for 9464D-GD2 tumors[18,19], after which treatments with individual or combination NAT components were given. Doses of treatment components for M2h include 1 µg of TNFα, 100 µg of aCD40, and 50 µg of Cetuximab. Doses of treatment components for 9464D-GD2 include 1 µg of TNFα, 100 µg of aCD40, and 50 µg of anti-GD2 mAb (clone 14G2a). Treatments were delivered intratumorally on the day of randomization, and a second dose was given 48 h later (**Fig. 1A**). Experiments involving neutrophil depletion used 500 µg of anti-Ly6G monoclonal antibody (BioXcell), administered intraperitoneally every other day for a total of five doses, beginning two days prior to the first treatment dose. Experiments involving ICI (250 µg of anti-PD1 and 200 µg of anti-CTLA4 antibodies) comprised of four doses delivered every seven days via intraperitoneal injections, starting seven days after the second dose of NAT in combination NAT + ICI treatment groups. All procedures were approved by the University of Wisconsin IACUC.

**Figure 1.**
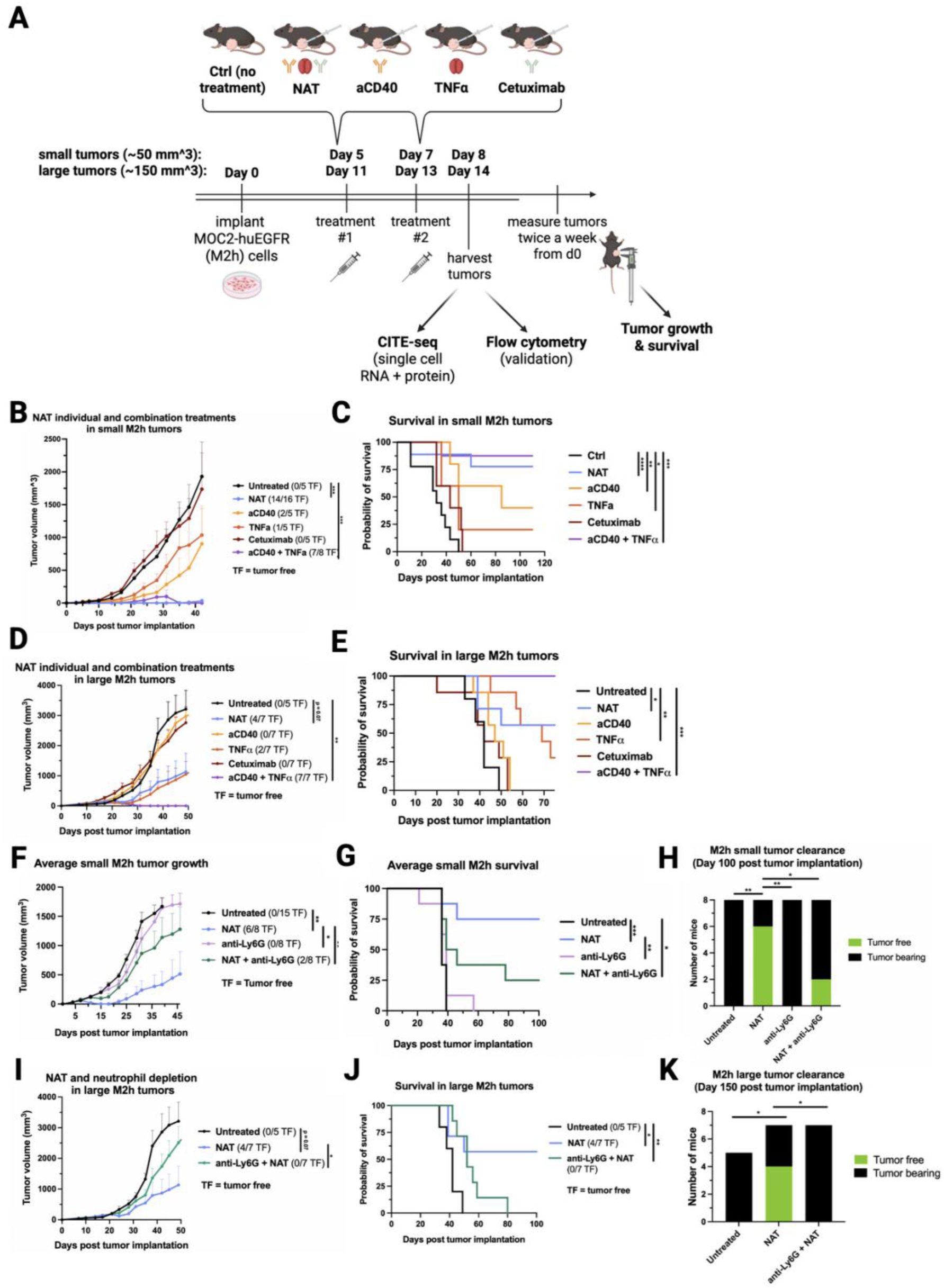
NAT improves M2h tumor control and clearance in part through neutrophil-dependent effects. (A) M2h tumor cells were implanted in mice and, once tumors reached small or large predefined sizes, animals were randomized to control (untreated), NAT, aCD40, TNFα, or Cetuximab treatment groups. Tumor growth, clearance, survival, and post-treatment immune phenotypes were then assessed by measuring tumor volume and analyzing harvested tumors with scRNA-seq, CITE-seq, and flow cytometry. (B-E) Mice bearing small or large M2h tumors were treated with NAT, aCD40, TNFα, Cetuximab, or aCD40 + TNFα, and tumor growth and overall survival were monitored relative to untreated controls. (F-H) In small M2h tumors, NAT alone or combined with anti-Ly6G was evaluated for effects on tumor growth, survival, and tumor clearance at day 100 post implantation. (I-K) In large M2h tumors, NAT alone or combined with anti-Ly6G was assessed for tumor growth, survival, and tumor clearance at day 150 post implantation. For all statistical comparisons, p<0.05 = *\**; p<0.01 *= **;* p<0.005 *= **\*\*\****; p<0.001 = ****.

### Single-cell multi-omics experiments

Large and small M2h tumors as well as small 9464D-GD2 tumors were collected 24 hours after the 2nd treatment dose, dissociated using the Miltenyi Mouse Tumor Dissociation Kit, and enriched for CD45^+^ cells (STEMCELL EasySep). For CITE-seq (large M2h tumors; 2–3 pooled/group), surface proteins were labeled with TotalSeq-C antibody-oligo conjugates (BioLegend); libraries were prepared per 10X Genomics Fixed RNA Profiling protocol (CG000673). For scRNA-seq (small M2h and 9464D-GD2 tumors), singleplexed fixed RNA profiling was used (CG000691). Libraries were sequenced on Illumina NovaSeq X Plus. Raw reads were processed with CellRanger (v8–9); downstream analysis was performed in Seurat (v4.3.0) with Harmony batch correction.[20] Cell types were annotated using canonical markers; doublets were removed using DoubletFinder or ScDblFinder.[21,22] Differentially expressed genes (DEGs) were identified using MAST (Bonferroni-adjusted FDR; log2FC cutoff 0.25).[23] Gene set enrichment analysis (GSEA) of Gene Ontology (GO) Biological Process terms, with Benjamini-Hochberg correction, was performed using clusterProfiler (v4.6.2) [24]. Cell-cell interactions were inferred using CellChat (v2.2.0).[25] Trajectory inferences were computed using Slingshot.[26] Large M2h tumor-derived N0-N4 neutrophil subpopulation annotations were transferred to small M2h tumor neutrophils and small 9464D-GD2 data by label transfer.[27]

### Flow cytometry

Large M2h tumors were dissociated, dead-cell depleted (Miltenyi), and CD45+ cells enriched (STEMCELL EasySep). Cells were stained with Ghost Red 780 (live/dead) and surface antibodies for CD11b, Ly6G, Ly6C, ICAM1 (CD54), DcTRAIL-R1, CCR3, and CD49d (Supplemental Table 1). FMO controls guided gating. Data were acquired on the Attune NxT machine and analyzed in FlowJo Version 10.10.1 Statistical comparisons used one-way ANOVA with Brown-Forsythe and Welch’s corrections (GraphPad Prism v10.6.1).

### HNSCC clinical data analysis

Neutrophils (CD11b+CD15+) were isolated from our previously published single-cell protein spatial CosMX SMI data[28] with 68 Tumor Microarray (TMA) cores on 23 HNSCC patients treated with ICIs (pembrolizumab-based therapy, either as a single agent or in combination) for abundance and ICAM1/CD54 expression analyses, with the Wilcox rank-sum test used to assess statistical significance. HNSCC overall survival analysis was performed using Kaplan-Meier survival analysis, with 500 HNSCC patients split by median ICAM1 expression, and ICAM1 together with human neutrophil markers (CSF3R, S100A8/9, NAMPT).

### Statistical methods

Durable tumor clearance is defined as complete elimination of all measurable/palpable tumor with no evidence of recurrence for at least 70 days (or as far out as the last day of the survival curve for that experiment). In figures, “tumor-free (TF)” represents animals with durable tumor clearance. For statistical analyses of tumor volume, the time-weighted average (area under the volume-time curve, calculated using trapezoidal method) was calculated for each mouse tumor. Time-weighted averages were compared between treatment groups overall by a Kruskal-Wallis test. If the overall test was found significant, then pairwise Mann-Whitney-Wilcoxon tests were conducted. Survival curves were compared with a log rank test overall, and then pairwise if significant. Tumor-free rates were compared with chi-square tests overall, and then pairwise proportion tests if significant. No p-value corrections were made for inflated type 1 error rate. Analysis was conducted using R version 4.5.2. Significance was assessed at the α = 0.05 level, and no adjustments were made to account for inflated type 1 error rate. Flow cytometry results are represented as scatter dot plots showing median (line) ± standard error of the mean. Each dot represents a single mouse tumor. A one-way ANOVA was calculated with Brown-Forsythe and Welch’s corrections, α = 0.05 (GraphPad Prism v10.6.1).

## RESULTS

### aCD40 and TNFα combination achieves comparable HNSCC tumor clearance as the full NAT regimen

To evaluate anti-tumoral efficacy of neutrophil-modulating therapies in an immunologically cold HNSCC context, we used the M2h model, which recapitulates key features of human HNSCC: a myeloid-rich, neutrophil-dominated TME with EGFR-expressed tumors, and resistance to ICI.[16] Tumor-bearing mice were treated with intratumoral NAT (aCD40 + TNFα + Cetuximab), its individual components, or aCD40 + TNFα, beginning at both small and large tumor sizes (**Fig. 1A**). A striking observation emerged with TNFα-containing regimens: after a single intratumoral dose of any such treatment in M2h tumors, extensive necrotic patches rapidly formed across most of the tumor surface, with aCD40 + TNFα or full NAT producing larger patches than TNFα monotherapy (**Suppl Fig. 1**). Across repeated experiments, these necrotic regions fell off over the course of two weeks, resulting in complete tumor clearance in most mice receiving double- or triple-agent therapy; by contrast, few TNFα monotherapy-treated mice with patches achieved clearance.

In both small and large M2h tumors, NAT achieved statistically significant, durable tumor clearance and improved survival compared with untreated controls (**Fig. 1B-E**). In contrast to the results with NAT by Linde et al.[12], in the M2h tumor aCD40+TNFα without Cetuximab was as effective as the full NAT regimen. For individual components, aCD40 monotherapy showed partial activity with mixed responses; TNFα produced delayed tumor growth without clearance; and Cetuximab alone was no more effective than untreated controls in both tumor growth inhibition and survival. The addition of ICI did not substantially improve the efficacy of either aCD40 monotherapy or NAT in small and large M2h tumors (**Suppl Fig. 2A-F**). Mice cured of small or large M2h tumors that remained tumor-free ∼70 days post-implant rejected tumor rechallenges, indicating durable memory (**Suppl Fig. 2G,H**).

To test the contribution of neutrophils to NAT-mediated tumor control, mice were depleted of neutrophils during NAT treatment, which significantly impaired NAT efficacy (**Fig. 1F-K**). These findings establish neutrophils as required mediators of the anti-tumoral response in this model and support the rationale for studying immunotherapy-induced reprogramming.

### Single-cell multi-omics identifies five transcriptionally distinct neutrophil states in M2h tumors following myeloid-modulating therapies

To characterize treatment-induced neutrophil heterogeneity, we performed CITE-seq on large M2h tumors, which showed shifts in immune cell frequencies, including B cells and macrophages but not neutrophils (**Fig. 2A-C**). As Cetuximab alone showed no anti-tumoral efficacy, this group was excluded from comparisons. Clustering identified five transcriptionally distinct neutrophil states: N0, N1, N2, N3, and N4, with markedly different frequencies across treatment conditions (**Fig. 2D-F**). N0, the dominant population in untreated tumors (89% of TANs), is characterized by matrix-remodeling (*Il1rap*, *Fgl2*, *Tgfbi*), T cell suppression via PD-L1 signaling (*Cmtm6*), and low ICAM1 RNA and protein expression, consistent with a tumor-promoting baseline state (**Fig. 2G,H**).[29] N0 frequency was markedly reduced across treatment conditions [aCD40 (7%), NAT (11%), TNFα (54%)] (**Fig. 2E,F**). N1 exhibited a pronounced type I interferon (IFN)-stimulated gene signature (*Ifi207*, *Ifit3*, *Gbp4/5/9, Gvin1*, *Xaf1*), with the highest ICAM1 levels of any cluster, recapitulating the recently reported ISG^+^ phenotypes[9,10] and showing markers of inflammasome (*Nod1*) and immunoproteasome (*Psme2b*) activity (**Fig. 2G,H**). N1 was rare in control tumors but expanded dramatically with aCD40 and NAT treatments, and to a lesser extent with TNFα (**Fig. 2 E,F**). N2 expressed hypoxia and stress-response adaptation (*Bnip3*, *Higd1a*, *Egln1*), metabolic reprogramming (*Hk2*, *Pfkp*, *Vegfa*), and DcTRAIL-R1 (*Tnfrsf23*) surface expression, with intermediate ICAM1 expression (**Fig. 2G**); N2 was most abundant with TNFα and NAT treatments, closely resembling previously described pro-tumoral T3 neutrophil states and *Ccl3*+ aged neutrophils (**Fig. 2G**).[13,38] N3 neutrophils expressed antimicrobial and effector genes (*Pglyrp1*, *Mmp8*, *Chil3*) at RNA level alongside CD193 (CCR3) at the surface protein level, with intermediate ICAM1 (**Fig. 2G,H**); this N3 population was nearly absent in untreated controls and aCD40-only conditions but expanded specifically with NAT and, to a lesser extent, with TNFα (**Fig. 2E,F**). We also detected N4, a small neutrophil precursor subset (CD49d/Itga4+) expressing antigen-presenting markers, present in all treatment groups (**Fig. 2E,F**). Integrated scRNA-Seq label transfer analysis of big and small tumors confirmed the induction of N1 in aCD40 and NAT, as well as enrichment of N0 in control and tumors treated with Cetuximab (**Suppl. Fig. 3A-C**). The N3 neutrophil subset was most abundant in small tumors treated with TNFα, but was also observed in other conditions, including control tumors (**Suppl Fig. 3A-C).** Overall, treated tumors exhibited greater neutrophil heterogeneity at the transcriptional and protein levels than controls, indicating distinct functional neutrophil states induced by different immunotherapies.

**Figure 2.**
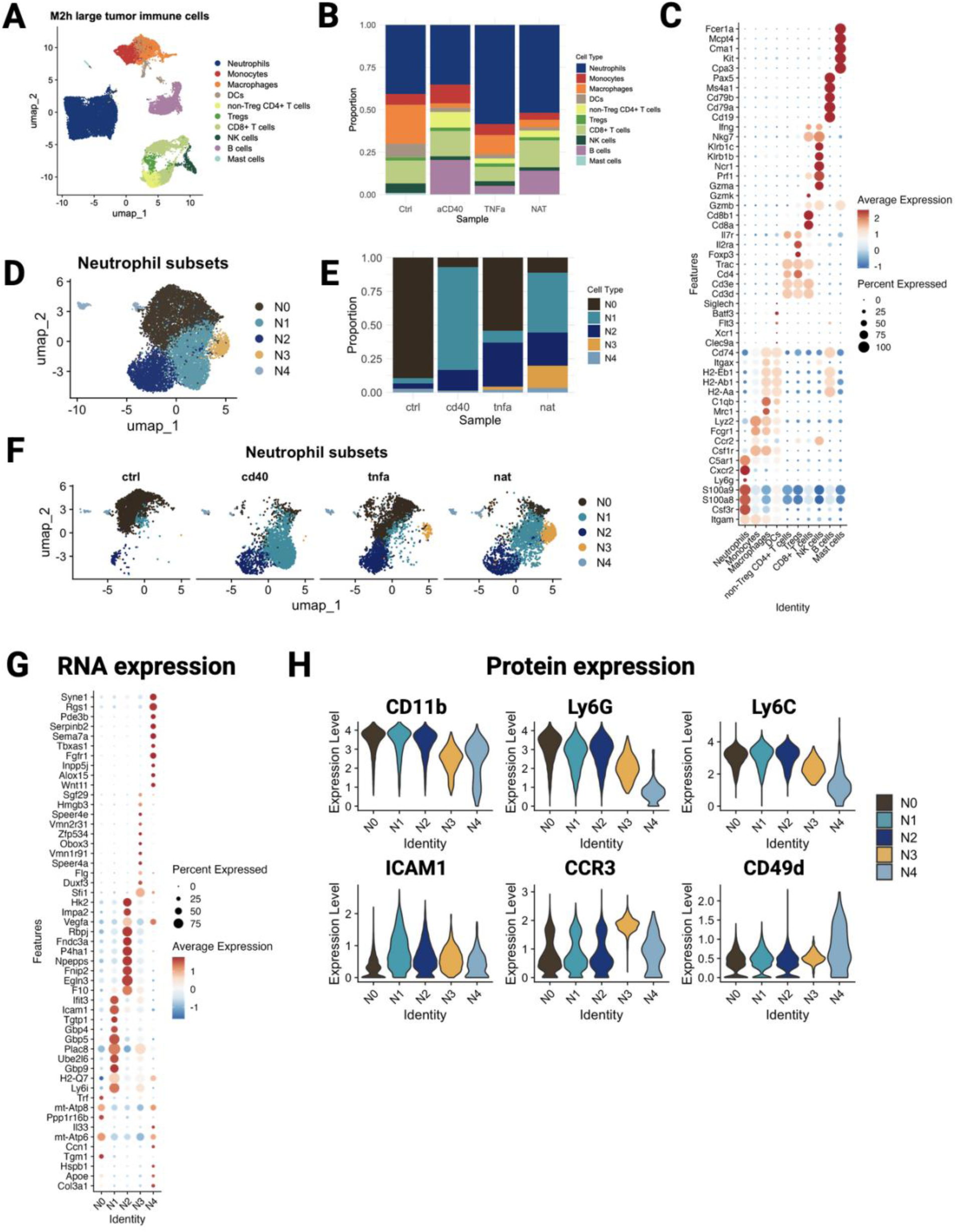
CITE-seq reveals treatment-associated remodeling of immune cells and identifies transcriptionally distinct neutrophil states in the large M2h tumor microenvironment. (A) UMAP of immune cells isolated from large M2h tumors, colored by annotated cell type. (B) Stacked bar chart showing the proportion of each immune cell type identified in (A) across treatment groups. (C) Dot plot of canonical marker genes used to annotate the immune cell clusters shown in (A), with dot size indicating the percentage of cells expressing each gene and color indicating average expression level. (D) UMAP of neutrophils isolated *in silico* from (A) and subclustered into five neutrophil states (N0-N4), colored by cluster. (E) Stacked bar chart showing the proportion of each neutrophil subset identified in (D) across treatment groups. (F) UMAPs of neutrophil subsets shown in (D), split by treatment group to visualize treatment-associated shifts in neutrophil state composition. (G) Dot plot of differentially expressed genes across neutrophil subsets (N0-N4), with dot size indicating the percentage of cells expressing each gene and color indicating average expression level. (H) Violin plots showing surface protein expression of CD11b, Ly6G, Ly6C, ICAM1, CCR3, and CD49d across neutrophil subsets (N0-N4), measured by CITE-seq antibody-derived tags.

### Flow cytometry validates treatment-induced neutrophil phenotypes

To confirm distinct neutrophil states at the protein level and establish a potential clinically applicable immunophenotyping assay, we designed and validated a flow cytometry panel using CD54(ICAM1), DcTRAIL-R1, CD193(CCR3), and CD49d as discriminating markers of neutrophil subpopulations (**Fig. 3A**). This panel was guided by CITE-seq surface protein data: N0 was gated as CD54-; N1 as CD54^+^CD193^-^DcTRAIL-R1^-^; N2 as CD54^+^DcTRAIL-R1^+^; N3 as CD54^+^CD193^+^; and N4 as CD49d^+^ (**Suppl. Fig. 4**). Neutrophil states identified with flow cytometry in large M2h tumors (**Fig. 3B**) confirmed that N0 neutrophils dominated untreated and Cetuximab-treated tumors and was significantly reduced across all other treatment conditions. N1 neutrophils increased significantly in all treatment groups versus untreated, except for remaining low in Cetuximab-treated tumors. N2 neutrophils expanded most significantly in aCD40 + TNFα and NAT groups relative to untreated controls. N3 neutrophils showed modest but statistically significant increases in aCD40+TNFα- and NAT-treated tumors versus controls and Cetuximab-treated tumors. N4 neutrophils remained <5% across all conditions, consistent with CITE-seq data. Quantitative differences between the flow and CITE-seq datasets, particularly the lower N1 magnitude in flow cytometry data, likely reflect biological and technical variables, including mouse sex, tumor implantation passage, and vivarium location, that differed between experimental cohorts.

**Figure 3.**
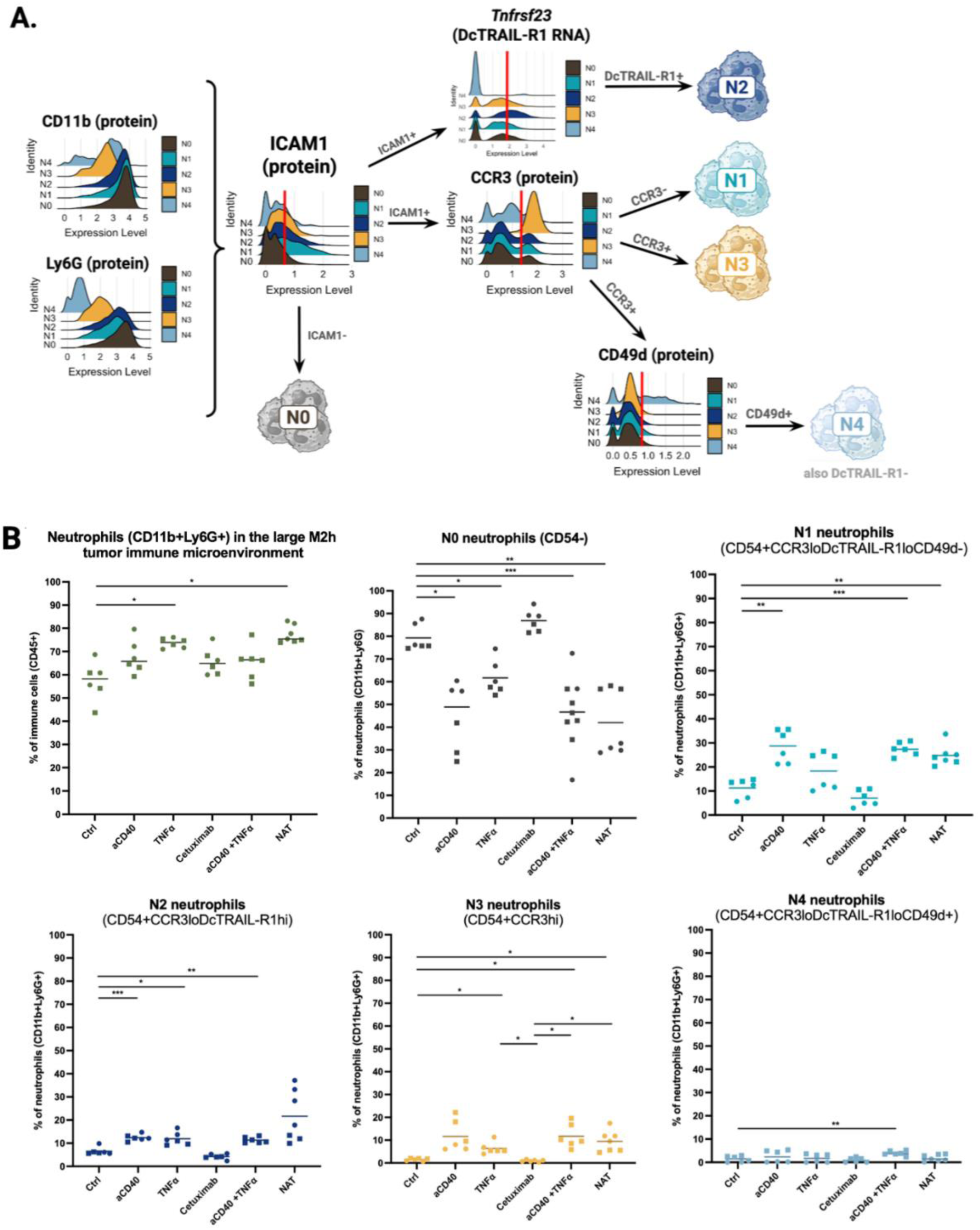
Validation of transcriptionally distinct neutrophil states in large M2h tumors by flow cytometry. (A) Schematic of the flow cytometry gating strategy used to identify neutrophil subsets (N0-N4) in large M2h tumors based on protein and RNA markers defined by CITE-seq. CD11b and Ly6G were used to identify neutrophils, CD54 separated N0 from CD54+ cells, and DcTrail-R1 (*Tnfrsf23*), CCR3, and CD49d were used to distinguish the remaining subsets. (B) Percentage of CD11b+Ly6G+ neutrophils among CD45+ immune cells across treatment groups. (C) Percentage of each neutrophil state (N0-N4) among total CD11b+Ly6G+ neutrophils across treatment groups. Flow cytometry was performed on large M2h tumors from two independent experiments. Each symbol represents an individual mouse tumor; circles and squares denote the two experiments. Statistical significance was assessed using one-way ANOVA with Welch and Brown-Forsythe correction followed by Dunnett’s T3 post-hoc test. For all statistical comparisons, p<0.05 = *\**; p<0.01 *= **;* p<0.005 *= **\*\*\****; p<0.001 = ****.

### Bioinformatics analyses predict distinct functions in treatment-induced neutrophil states

To explore the functional context across different neutrophil states, we performed GSEA using GO biological process terms and KEGG metabolic and apoptosis pathways for each neutrophil cluster (**Fig. 4A, Suppl. Fig. 5A,B**). N0 showed sparse pathway enrichment overall, with active suppression of innate immune activation pathways, consistent with a functionally restrained state (**Fig. 4A**). Average expression comparison across representative KEGG pathways supported metabolic quiescence features in N0, with lower expression of gene signatures for glycolysis, oxidative phosphorylation, and the TCA cycle (**Suppl. Fig. 5A**). N1 and N3 statistically shared enrichment across a broad panel of innate immune effector pathways, including phagocytosis, defense response to bacterium, IFN signaling, leukocyte-mediated immunity, pattern recognition receptor signaling, with only N3 gene signatures associated with regulation of actin dynamics and cell migration (leukocyte chemotaxis) (**Fig. 4A**). N2 neutrophils did not enrich for any GO biological process pathways after multiple testing correction (**Fig. 4A**). N3 was the most metabolically active cluster across nearly all KEGG pathways (**Suppl. Fig. 5A**), combined with FDR-corrected enrichment in leukocyte chemotaxis (the most gene-dense and statistically robust enrichment in the full dataset) and antimicrobial defense, alongside statistically robust suppression of MHC complex assembly and antigen presentation (**Fig. 4A**). N4 showed exclusively developmental and vascular gene signatures, profound metabolic suppression, and the strongest apoptotic resistance of any cluster, confirming its identity as a metabolically dormant, functionally inert progenitor-like state (**Suppl. Fig. 5A,B**).

**Figure 4.**
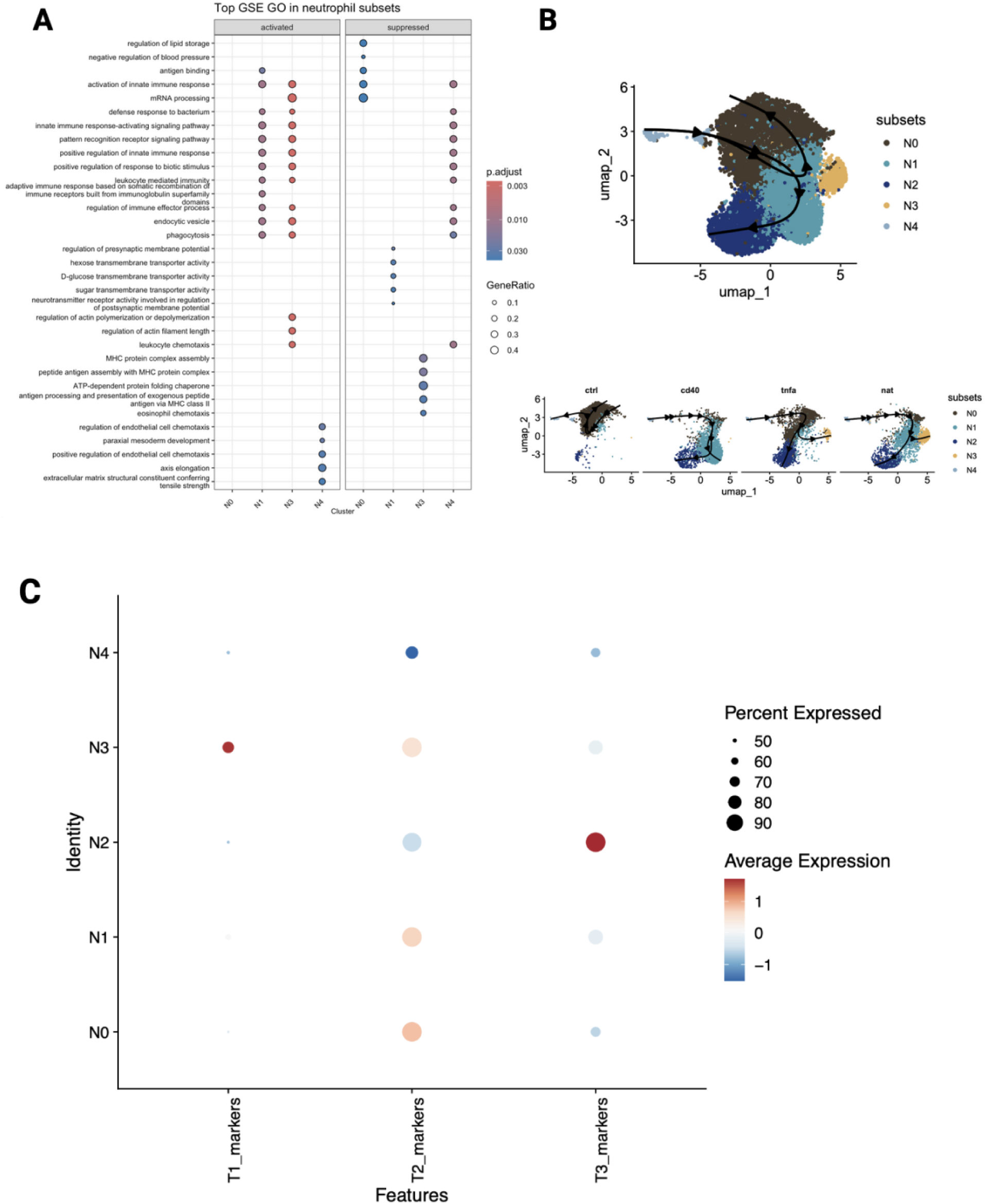
Complementary pathway, trajectory, and signature analyses further define neutrophil state predictions in large M2h tumors. (A) Dot plot of top gene set enrichment Gene Ontology terms activated and suppressed across neutrophil subsets (N0-N4), with dot size indicating gene ratio and color indicating adjusted p-value. (B) UMAP of large M2h neutrophil subsets (N0-N4) overlaid with predicted developmental trajectories inferred by Slingshot analysis, with arrows indicating the direction of state transitions across all treatment groups combined; below, the same UMAPs are split by treatment group to show treatment-associated differences in predicted neutrophil state transitions. (C) Dot plot of previously defined T1, T2, and T3 neutrophil state gene signatures across neutrophil subsets (N0-N4), with dot size indicating the percentage of cells expressing each signature and color indicating average expression level. N1 was most concordant with T2, N3 with T1, and N2 with T3, supporting the trajectory analysis shown in (B).

Slingshot trajectory analysis predicted treatment-dependent neutrophil state paths, all originating from the progenitor-like N4 state (**Fig. 4B**). Overall, the bioinformatics results suggested distinct potential bifurcated neutrophil induction in TME with a shared established terminal differentiated state, N2 across all treatments, similar to ones identified in the literature[7,8]. However, the alternative trajectories toward anti-tumoral states, including aCD40 with ISG^+^ N1 states (% of N1 compared to N2), TNFɑ with CCR3^+^ N3 states, and NAT with both N1+N3 states, suggest potential resistance to becoming terminal pro-tumoral N2 states or newly recruited states. Using T1-T3 neutrophil state gene signatures[7], we found enrichment of N1 in T2 and N3 in T1 states, while N2 is highly concordant with T3 states (**Fig. 4C**), supporting our trajectory analysis.

### Treatment-induced neutrophils associate with T cell intercellular communication

Using T cell phenotype as a readout of anti-tumor neutrophil feature in the absence of sufficient tumor cells following treatment, we used CellChat analysis to look at Ligand-Receptor (LR) interactions. Treatment conditions suggested a significant increase in both the number and strength of LR interactions compared to untreated tumors, with TNFα and NAT producing the most expansive changes, particularly with CD8+ T cells as receiver cells (**Fig. 5B**). While ISG^+^ N1 has the highest outgoing signaling overall, N3 has the highest in NAT, suggesting the potentially important synergy of this migration-associated neutrophil state. N3 was absent from untreated tumors and aCD40, indicating it as a strictly treatment-emergent population associated with TNFα (**Fig. 5B** and **Fig. 2F,G**). N2 has overall lower LR interaction enrichment than N1 in all and N3 in TNFα-associated treatment, indicating that its interactions with T cells in pro-tumoral states are not outnumbered by those in anti-tumoral states in successful treatments. Non-Treg CD4+ T cells and Tregs maintained relatively consistent communication patterns across conditions, suggesting their signaling roles were less sensitive to these specific interventions (**Fig. 5B**).

**Figure 5.**
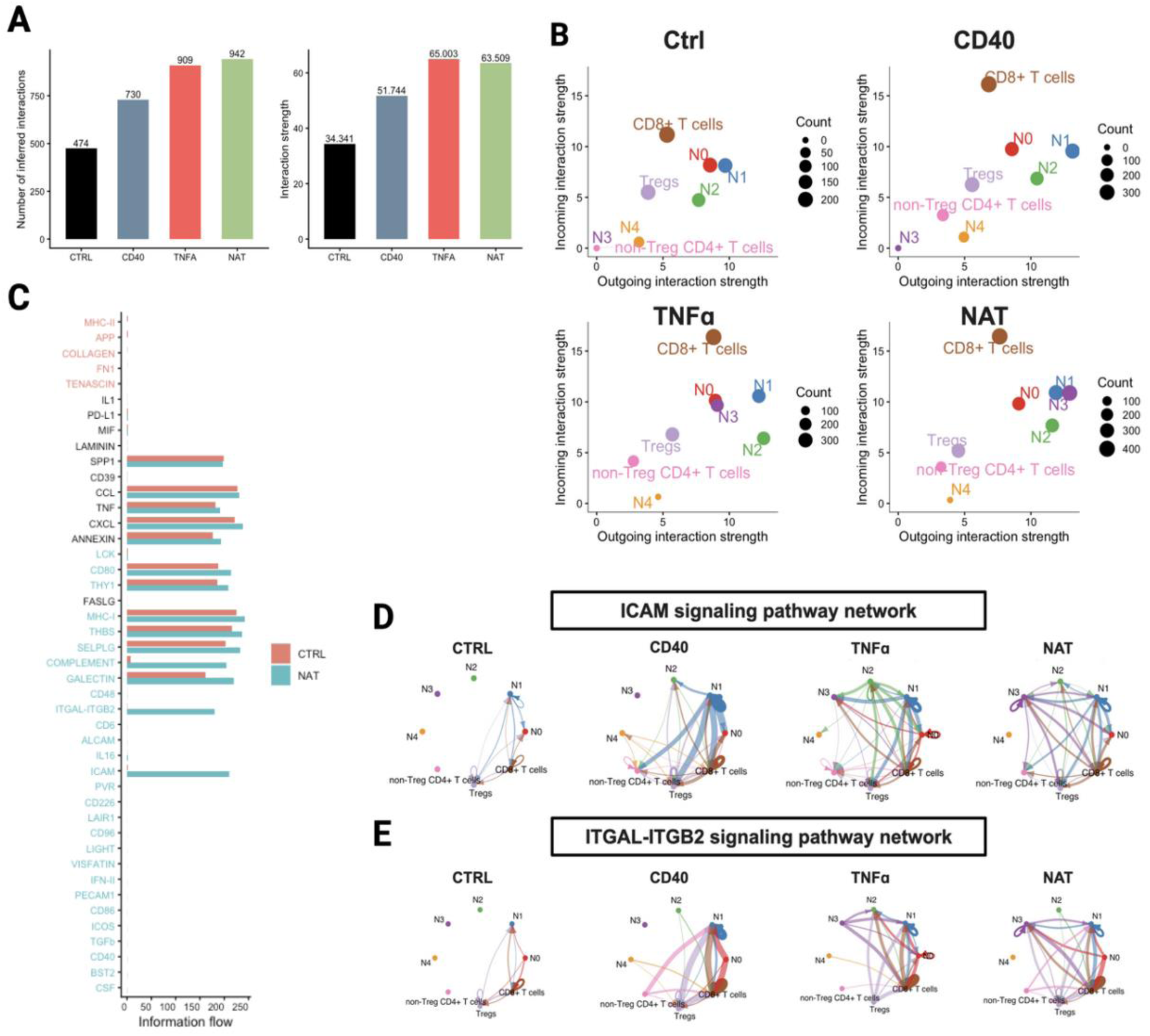
CellChat ligand-receptor analyses reveal treatment-induced remodeling of neutrophil-T cell communication networks in large M2h tumors. (A) CellChat was applied to the large M2h tumor CITE-seq data to infer intercellular communication across treatment groups. Total inferred interactions and overall interaction strength across control (untreated), aCD40, TNFα, and NAT treatment groups. (B) Outgoing versus incoming interaction strength for each cell type are shown, with dot size indicating the number of inferred interactions. (C). Relative and total information flow for each signaling pathway in untreated control and NAT groups. (D) and (E) ICAM and ITGAL-ITGB2 signaling chord diagrams among neutrophil subsets and T cells across treatment groups are shown, with arrow color indicating signal origin and arrow width indicating signal strength.

Comparison of signaling pathways between untreated and NAT conditions suggested changes in the TME communication architecture with COMPLEMENT, ITGAL-ITGB2, and ICAM pathways as highly enriched in NAT-treated tumors (**Fig. 5C**). Examination of the ICAM signaling network across conditions showed that in untreated tumors, ICAM interactions were mostly between N0, N1, and CD8+ T cells while ISG^+^ N1 emerged as a dominant hub with a prominent autocrine loop and strong outgoing signals to CD8+ T cells for aCD40-treated tumors (**Fig. 5D**). For TNFα treated tumors the network became the most broadly connected, with N3 engaging for the first time in bidirectional ICAM signaling. With NAT treatment, N1 and N3 were reinforced as both senders and receivers, with persistent autocrine ICAM signaling in N1 and CD8+ T cells (**Fig. 5D**). Similar results were observed in the ITGAL-ITGB2 (LFA-1) network, a key integrin pathway regulating neutrophil-T cell adhesion and immunological synapse formation (**Fig. 5E**).The consistent emergence of N1 as a central hub across both networks, and the NAT-specific enhancement of N3 engagement over that of TNFα alone, identifies these adhesion and co-stimulatory axes as likely core mechanisms through which treatment-induced neutrophils communicate with CD8+ T cells.

### NAT induces new neutrophil states across tumor models

To assess the generalizability of NAT-induced neutrophil state shifts, we tested NAT (aCD40 + TNFα + anti-GD2 mAb) in the syngeneic 9464D-GD2 neuroblastoma model. NAT significantly prolonged survival compared to untreated controls and achieved durable tumor clearance in a small subset of mice (**Fig. 6A-C**). While this shows that NAT has significant anti-tumor activity against the 9464D-GD2 tumor, its anti-tumor potency is substantially weaker when the same dose and regimen are used in the same mouse strain against the syngeneic M2h tumor of similar size (**Fig. 1B, C**). Notably, neutrophil depletion did not significantly impair NAT efficacy in 9464D-GD2, in contrast to the neutrophil dependence observed for NAT efficacy in M2h tumors, indicating that that the mechanistic contribution of neutrophils to NAT-mediated tumor control is tumor-type specific (**Fig. 6A-C**).

**Figure 6.**
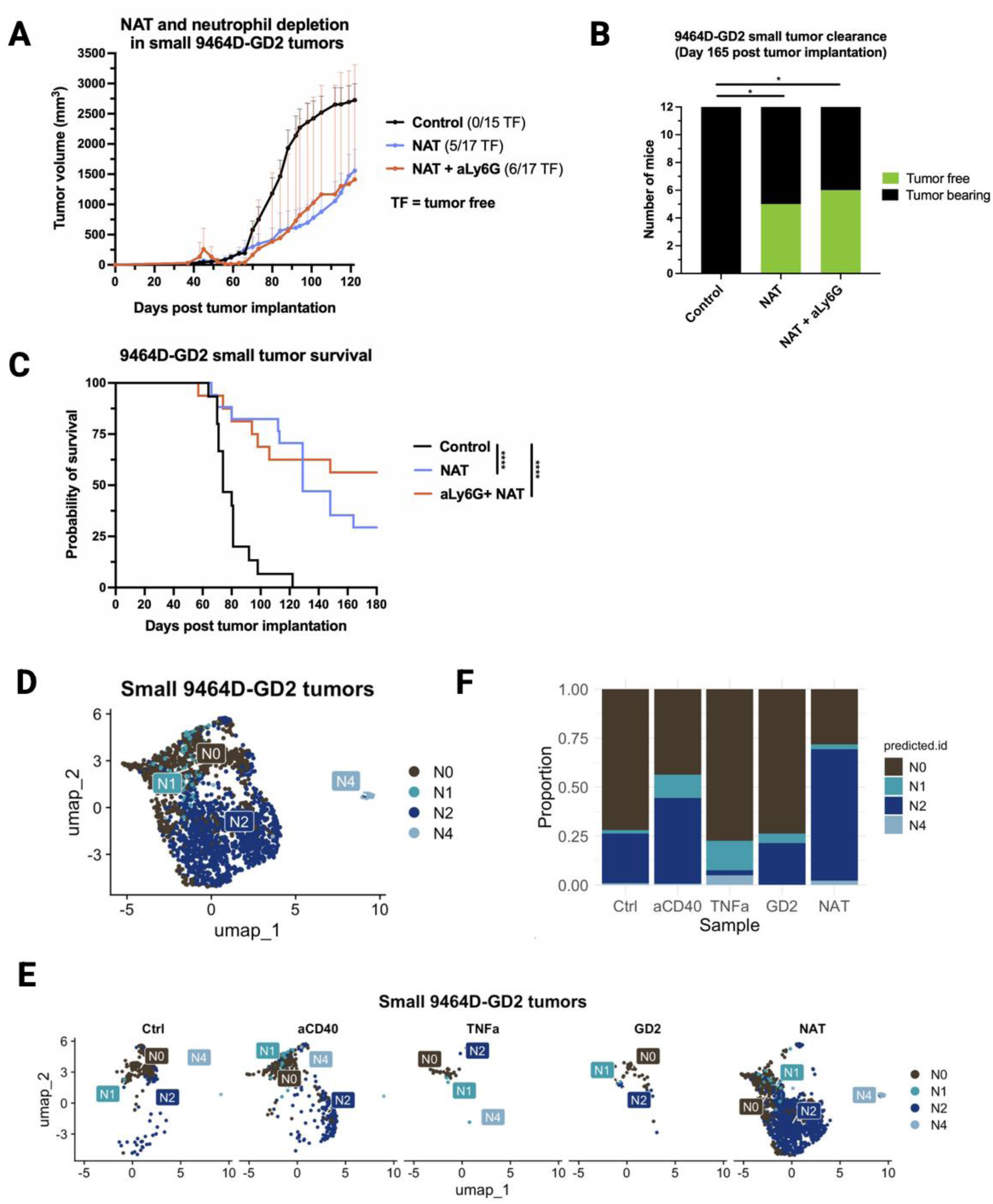
NAT prolongs survival and induces partial tumor clearance in small 9464D-GD2 tumors, while label transfer reveals partial conservation of treatment-associated neutrophil state shifts across tumor models. (A) Average tumor volume (+ SEM) of mice bearing small 9464D-GD2 flank tumors left untreated (control), treated with NAT, or treated with NAT + aLy6G (n = 15-17/group). TF = tumor free. (B) Number of tumor-bearing and tumor-free mice in each group at day 165 post tumor implantation. (C) Overall survival of mice in each treatment group. For all statistical comparisons, p<0.05 = *\**; p<0.01 *= **;* p<0.005 *= \*\*\**; p<0.001 = ****. (D)Neutrophils were subsetted *in silico* from the small 9464D-GD2 scRNA-seq dataset and assigned subset identities by label transfer using the large M2h tumor neutrophil annotations (N0-N4) as a reference. UMAP of small 9464D-GD2 tumor-infiltrated neutrophils colored by label transfer-predicted subset identity. (E) UMAPs as in (D), split by treatment group to visualize treatment-associated shifts in neutrophil subset composition. (F) Stacked bar chart showing the proportions of each predicted neutrophil subset across treatment groups.

We then performed scRNA-seq of small 9464D-GD2 tumors treated with individual and combination components of NAT. Computational label transfer of M2h-derived neutrophil states onto the scRNA-seq data for 9464D-GD2 neutrophils identified all but N3 states (**Fig. 6D-F**). First, N0 dominated untreated and single-agent-treated 9464D-GD2 tumors, mirroring its dominance in untreated M2h tumors, a cross-model conservation that reinforces N0 as a pro-tumoral, immunosuppressive TAN state in cold tumors (**Fig. 6E,F**). Second, NAT was the only condition that substantially reduced N0 frequency in both models, confirming that NAT, in synergy with aCD40 and TNFɑ, has a conserved capacity to dismantle the baseline immunosuppressive neutrophil landscape across tumor types. However, none of the treatments induced sufficient ISG^+^ N1 states relative to hypoxia-associated N2, which may help explain why neutrophil depletion improved outcomes in this model (**Fig. 6C**). Third, an NAT-expanded neutrophil cluster in 9464D-GD2 characterized by *Nos2* expression, hypoxia-response genes, and glycolytic reprogramming mapped predominantly to the M2h N2 identity rather than N1 (scRNAseq data not shown), introducing ambiguity as to whether this population is anti-tumoral or represents a stress-adapted state. N4 was conserved at consistently low frequencies across all conditions in both models. Finally, N3 was essentially absent from 9464D-GD2 tumors (and thus not shown in **Fig.6 D-F)**, suggesting that the full repertoire of NAT-induced TAN states observed in M2h is not universally reproduced across tumor types (**Fig. 6E,F**), and that neutrophil-activating mechanisms are likely differ between these two models

### CD54(ICAM1) expression in neutrophils associates with ICI response and survival in HNSCC patients

Because the clinical use of aCD40 combined with TNFα in HNSCC patients has not yet been tested, we used Pembrolizumab (anti-PD-1, a component of the ICI used in our murine studies above) treatment response as a clinically accessible surrogate for potential immunotherapy-induced anti-tumoral neutrophil activity. We reanalyzed a previously published CosMX SMI spatial protein dataset comprising 68 tumor microarray (TMA) cores from 23 HNSCC patients (10 responders, 13 non-responders) treated with pembrolizumab-based ICI (**Fig. 7**) [28]. ICAM1 protein expression on tumor-infiltrating neutrophils was higher in responders than non-responders at the TMA core level, though this trend did not reach conventional significance when analyzed with a single value for each of the 23 patients, most likely due to limited cohort size and inter-patient heterogeneity. Neutrophil abundance was comparable between responders and non-responders, suggesting that ICAM1 reflects a qualitative activation difference rather than a quantitative neutrophil infiltration difference (**Fig. 7A,B**).

**Figure 7.**
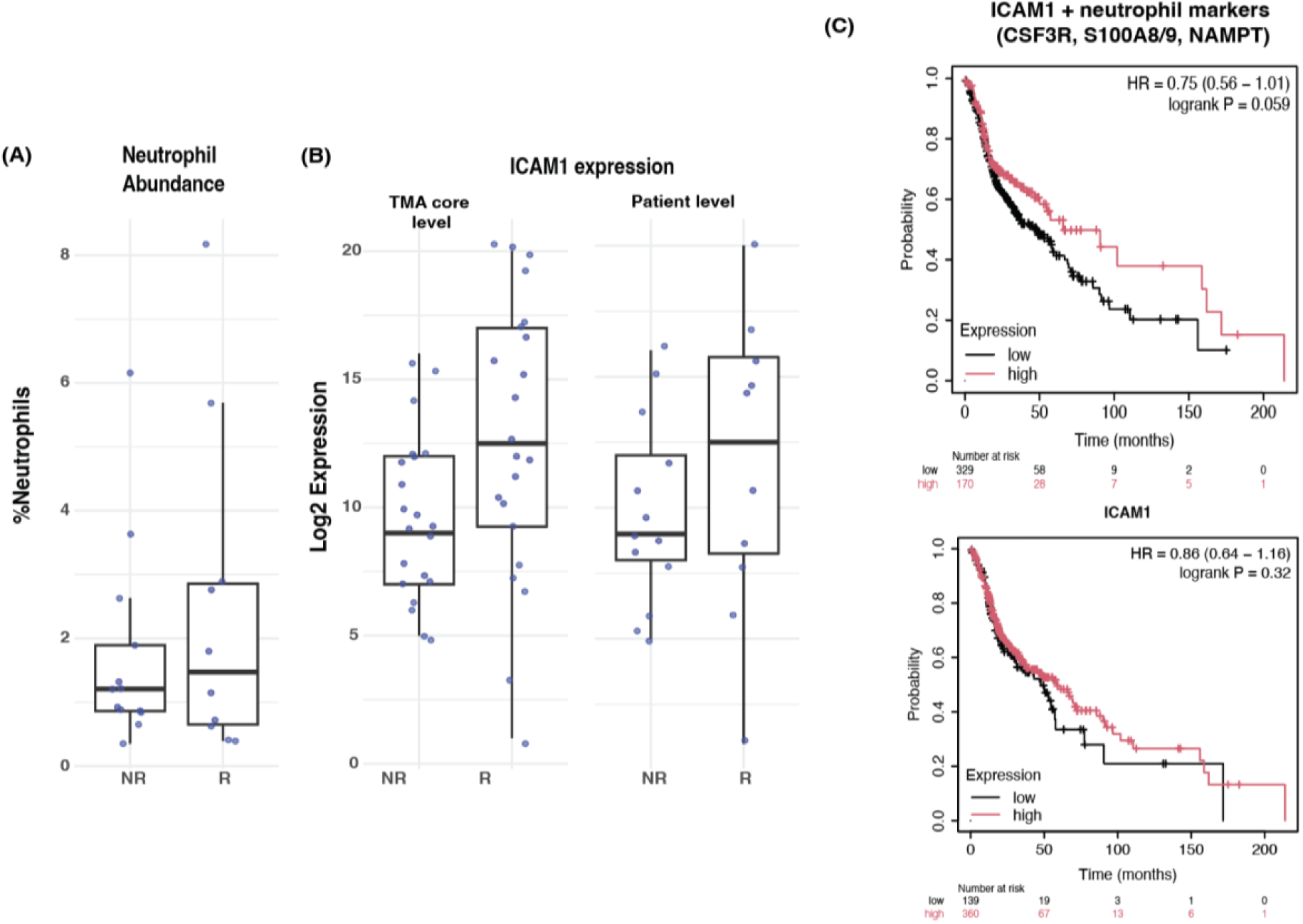
ICAM1^+^ neutrophils is a potential marker for anti-tumoral neutrophils. Evaluation of neutrophils in spatial data of HNSCC tumors from ICI Responders and Non-Responders with (A) neutrophil abundance, (B) ICAM1/CD54 log2 expression (n=13 Non-Responders, n= 10 Responders), and ICAM1/CD54 expression at the TMA core level (n=21 Non-Responders, n=22 Responders). (C) Kaplan-Meier survival analysis of HNSCC patients split by high and low from median expression of ICAM1 and neutrophil markers (CSF3R, S100A8/9, NAMPT, above) and ICAM1 alone (below).

To assess whether ICAM1+ neutrophil signatures are associated with patient survival more broadly, we performed a Kaplan-Meier analysis in the TCGA HNSCC cohort (n=500 patients). When patients were stratified by ICAM1 expression alone, no significant survival difference was detected (HR=0.86, p=0.32) (**Fig. 7C**). However, stratification by a composite score combining ICAM1 with canonical neutrophil identity markers (CSF3R, S100A8/9, NAMPT), which enriches specifically for ICAM1-expressing neutrophils rather than other ICAM1^+^ cell types, produced a trend toward improved overall survival (HR=0.75, p=0.059) (**Fig. 7C**). The directional consistency of these human data with the mouse model findings, across two independent patient cohorts, supports the translational potential of ICAM1 as a treatment-induced neutrophil activation biomarker and warrants prospective evaluation in larger patient cohorts.

## DISCUSSION AND CONCLUSIONS

This study presents a single-cell multi-omics framework for evaluating how different myeloid-modulating immunotherapies alter TAN phenotypes in HNSCC, a cancer type characterized by an immunologically cold, myeloid-dominated TME and poor responsiveness to ICI. Using integrated immunoprofiling and bioinformatics analyses in the M2h syngeneic model, we identify five transcriptionally and functionally distinct TAN states (N0–N4), validated by flow cytometry, and provide cross-model and translational evidence that CD54/ICAM1 marks treatment-activated neutrophils with anti-tumor signatures with potential clinical relevance. The central conceptual advance is that successful immunotherapies reprogram neutrophils not merely in their abundance but also in kind: the qualitative nature of reprogramming, not its magnitude, appears to distinguish effective from ineffective treatment. This perspective supports emerging reframes of neutrophils from undifferentiated immune suppressors into a heterogeneous and potentially therapeutically malleable cell population whose state composition could serve both as a mechanistic readout and as a biomarker of treatment response.

The unexpected comparability of aCD40 + TNFα to full NAT treatment for M2h tumors, not oberved in the B16 melanoma model presented in the 2023 Linde et al. NAT study, warrants consideration.[12] Thus, the requirement for an anti-tumor antibody in the NAT regimen appears to be tumor-dependent (e.g., B16), but not universal (e.g., M2h). Additional studies are needed to define the spectrum of murine tumors that require inclusion of an anti-tumor mAb to NAT for efficacy, versus those in whichaCD40 and TNFα are sufficient, and to identify tumor-intrinsic characteristics that determine this dependence. If aCD40 + TNFα can drive equivalent tumor clearance through N1-mediated innate effector mechanisms for a large number of tumors, the translational implications would be significant. A major limitation of antibody-based tumor-targeting is antigen heterogeneity and loss. A regimen that does not require a defined tumor antigen would be inherently resistant to this failure mode.

The neutrophil-dependence of NAT efficacy in M2h, established by anti-Ly6G depletion, contrasts with the dispensability of neutrophils in 9464D-GD2 and mirrors findings from Linde et al.[14] The ISG^+^ N1 state, characterized by high CD54/ICAM1 expression and other IFN-α/β-stimulated genes, is the dominant neutrophil state induced by aCD40 and remains highly abundant with NAT. Its enrichment in phagocytosis, defense response, IFN signaling, and leukocyte-mediated immunity pathways is consistent with an anti-tumoral functional state, in line with prior reports of ISG^+^ neutrophils as anti-tumoral effectors.[10] CellChat analysis of ICAM^-^ and LFA-1-mediated N1-CD8^+^ T cell interactions positions N1 neutrophils not only as intrinsic cytotoxic effectors, but also as an active bridge between innate and adaptive immunity in the treated TME. However, N1’s apoptotic priming signature, consistent with the short lifespan of mature neutrophils, suggests a temporally limited anti-tumoral effect. Strategies to extend N1 survival *in vivo*, such as G-CSF, survival niche-promoting signals, or anti-apoptotic co-treatments, may therefore be required to sustain its activity beyond the acute treatment window.

The expansion of the N3 state following NAT and TNFα is another mechanistically notable finding. N3 is absent from untreated tumors, indicating a strictly treatment-induced population. Its high ICAM1 expression and enrichment in leukocyte chemotaxis and antimicrobial effector pathways suggest a highly activated effector state. The unique surface expression of CD193/CCR3 on N3, detectable by CITE-seq antibody-oligo quantification but absent at the RNA level, suggests post-transcriptional or translational regulation and may indicate a role for CCR3 in intratumoral localization or retention. Given prior evidence that CCR3 can modulate neutrophil recruitment in inflammation settings[30], its functional role in tumors warrants further investigation.

We also developed and validated of a flow cytometry panel, including CD54/ICAM1 and CD193/CCR3, for future functional and clinically applicable assays. Prospective validation in larger patient cohorts, ideally with paired pre- and post-treatment tumor neutrophil profiling, will be required to establish ICAM1 neutrophils as a clinically important biomarker. The cross-model analysis in 9464D-GD2 neuroblastoma reinforces and refines our understanding of NAT-induced neutrophil states. The model-specific differences in which neutrophil states are induced likely depend on tumor-intrinsic factors. It may be that for M2h, depletion of neutrophils interferes with the anti-tumor effect because the treatment induces heterogeneous neutrophil states that have a net balance of anti-tumor activity. In contrast, for the 9464D-GD2 tumors, NAT induces a shift in the neutrophil state distribution, with a preponderance of N2 neutrophils, few N1, and no N3; this might indicate that post-treatment pro-tumoral neutrophils outweigh anti-tumoral neutrophils. These results caution against assuming a universal mechanism for NAT across tumor types and underscores the need for tumor-specific mechanistic characterization.

Several limitations warrant acknowledgment. Functional assignment of neutrophil states are inferred from transcriptional and protein single-cell rather than direct functional assays;*ex vivo* cytotoxicity assays, co-culture, and T cell activation assays will be required for validation. Computational trajectories, based on static single-cell snapshots cannot establish lineage relationships and should be complemented by time-resolved or lineage-tracing approaches, which are challenging for neutrophil studies. Finally, the human cohorts analyzed are limited by size and not fully matched to the therapeutic regimens modeled in mice, contraining statistical power and clinical generalizability.

Together, these findings support a model in which the balance between stress- or hypoxia-associated immunosuppressive/pro-tumoral and ISG-associated anti-tumoral states determines the net therapeutic outcome. Rational interventions that selectively expand anti-tumor neutrophil states while constraining pro-tumor/immunosuppressive populations may therefore enhance efficacy and represent a promising avenue for improving responses in myeloid-rich, immunologically cold tumors.

## Availability of data and material

Raw sequencing data will be deposite in GEO upon acceptance. All code used for data analysis is available upon reasonable request.

## Authors’ contributions

AG and HQD conceived and designed the study. AG carried out the bulk of the data collection, with help from SS, NW, AKE, SD, KH, SB, JM, and ST. AG and HQD performed computational analyses. JZ performed statistical analyses on tumor growth and survival experiments. AG, AKE, PMS, and HQD interpreted the data. AG drafted the manuscript, which was edited by all coauthors. All authors reviewed and approved the final version.

## Ethics approval

All animal experiments were conducted under protocols approved by the University of Wisconsin Institutional Animal Care and Use Committee (IACUC). Human patient data were derived from previously published and de-identified cohorts; no new human subjects research was conducted.

## Acknowledgements

We thank the McArdle Laboratory for Cancer Research, Department of Human Oncology, and the Carbone Cancer Center at the University of Wisconsin-Madison for their ongoing support. The authors thank the UWCCC Flow Laboratory, WIMR Vivarium staff, Translational Research Initiatives in Pathology, UWCCC Informatics Shared Resource and Biomedical Research Model Services for use of their facilities and services. Additionally, the authors thank Wei Wang, PhD, Paul F. Lambert, PhD, Rong Hu, MD, PhD, and Melissa Meyer, PhD (Des Moines University) for their guidance and insightful feedback.

## REFERENCES

1. Hedrick CC, Malanchi I. Neutrophils in cancer: heterogeneous and multifaceted. Nat Rev Immunol. 2022;22:173–87.

2. Masucci MT, Minopoli M, Carriero MV. Tumor Associated Neutrophils. Their role in tumorigenesis, metastasis, prognosis and therapy. Front Oncol. 2019;9:1146.

3. Hao Y, Hu P, Zhang J. Genomic analysis of the prognostic effect of tumor-associated neutrophil-related genes across 15 solid cancer types: an immune perspective. Ann Transl Med. 2020;8:1507.

4. Bekes EM, Schweighofer B, Kupriyanova TA, Zajac E, Ardi VC, Quigley JP, et al. Tumor-recruited neutrophils and neutrophil TIMP-free MMP-9 regulate coordinately the levels of tumor angiogenesis and efficiency of malignant cell intravasation. Am J Pathol. 2011;179:1455–70.

5. Spiegel A, Brooks MW, Houshyar S, Reinhardt F, Ardolino M, Fessler E, et al. Neutrophils suppress intraluminal NK cell-mediated tumor cell clearance and enhance extravasation of disseminated carcinoma cells. Cancer Discov. 2016;6:630–49.

6. Wu L, Saxena S, Awaji M, Singh RK. Tumor-associated neutrophils in cancer: Going pro. Cancers (Basel). 2019;11:E564.

7. Ng MSF, Kwok I, Tan L, Shi C, Cerezo-Wallis D, Tan Y, et al. Deterministic reprogramming of neutrophils within tumors. Science. 2024;383:eadf6493.

8. Bolli E, Wirapati P, Hicham M, Xie Y, Siwicki M, Duval F, et al. CCL3 is produced by aged neutrophils across cancers and promotes tumor growth. Cancer Cell [Internet]. 2026; Available from: 10.1016/j.ccell.2026.01.006

9. Gungabeesoon J, Gort-Freitas NA, Kiss M, Bolli E, Messemaker M, Siwicki M, et al. A neutrophil response linked to tumor control in immunotherapy. Cell. 2023;186:1448–1464.e20.

10. Benguigui M, Cooper TJ, Kalkar P, Schif-Zuck S, Halaban R, Bacchiocchi A, et al. Interferon-stimulated neutrophils as a predictor of immunotherapy response. Cancer Cell. 2024;42:253–265.e12.

11. Hirschhorn D, Budhu S, Kraehenbuehl L, Gigoux M, Schröder D, Chow A, et al. T cell immunotherapies engage neutrophils to eliminate tumor antigen escape variants. Cell. 2023;186:1432–1447.e17.

12. Linde IL, Prestwood TR, Qiu J, Pilarowski G, Linde MH, Zhang X, et al. Neutrophil-activating therapy for the treatment of cancer. Cancer Cell. 2023;41:356–372.e10.

13. Larkins E, Blumenthal GM, Yuan W, He K, Sridhara R, Subramaniam S, et al. FDA approval summary: Pembrolizumab for the treatment of recurrent or metastatic head and neck squamous cell carcinoma with disease progression on or after platinum-containing chemotherapy. Oncologist. 2017;22:873–8.

14. Luo R, Yang J, Cao Z, Li B. Myeloid-driven immunosuppression in head and neck cancer: single-cell ATAC/RNA and spatial transcriptomic perspectives. Front Oncol. 2025;15:1693152.

15. Kasahara Y, Saijo K, Ueta R, Numakura R, Sasaki K, Yoshida Y, et al. Pretreatment neutrophil-lymphocyte ratio as a prognostic factor in recurrent/metastatic head and neck cancer treated with pembrolizumab. Sci Rep. 2024;14:28255.

16. Wang W, Lozar T, Golfinos AE, Lee D, Gronski E, Ward-Shaw E, et al. Stress keratin 17 expression in head and neck cancer contributes to immune evasion and resistance to immune-checkpoint blockade. Clin Cancer Res. 2022;28:2953–68.

17. Kono M, Saito S, Egloff AM, Allen CT, Uppaluri R. The mouse oral carcinoma (MOC) model: A 10-year retrospective on model development and head and neck cancer investigations. Oral Oncol. 2022;132:106012.

18. Jin WJ, Erbe AK, Schwarz CN, Jaquish AA, Anderson BR, Sriramaneni RN, et al. Tumor-specific antibody, cetuximab, enhances the in situ vaccine effect of radiation in immunologically cold head and neck squamous cell carcinoma. Front Immunol. 2020;11:591139.

19. Voeller J, Erbe AK, Slowinski J, Rasmussen K, Carlson PM, Hoefges A, et al. Combined innate and adaptive immunotherapy overcomes resistance of immunologically cold syngeneic murine neuroblastoma to checkpoint inhibition. J Immunother Cancer. 2019;7:344.

20. Korsunsky I, Millard N, Fan J, Slowikowski K, Zhang F, Wei K, et al. Fast, sensitive and accurate integration of single-cell data with Harmony. Nat Methods. 2019;16:1289–96.

21. Germain P-L, Lun A, Macnair W, Robinson MD. Doublet identification in single-cell sequencing data using scDblFinder. F1000Res. 2021;10:979.

22. Germain P-L, Lun A, Garcia Meixide C, Macnair W, Robinson MD. Doublet identification in single-cell sequencing data using scDblFinder. F1000Res. 2021;10:979.

23. Finak G, McDavid A, Yajima M, Deng J, Gersuk V, Shalek AK, et al. MAST: a flexible statistical framework for assessing transcriptional changes and characterizing heterogeneity in single-cell RNA sequencing data. Genome Biol. 2015;16:278.

24. Wu T, Hu E, Xu S, Chen M, Guo P, Dai Z, et al. clusterProfiler 4.0: A universal enrichment tool for interpreting omics data. Innovation (Camb). 2021;2:100141.

25. Jin S, Guerrero-Juarez CF, Zhang L, Chang I, Ramos R, Kuan C-H, et al. Inference and analysis of cell-cell communication using CellChat. Nat Commun. 2021;12:1088.

26. Street K, Risso D, Fletcher RB, Das D, Ngai J, Yosef N, et al. Slingshot: cell lineage and pseudotime inference for single-cell transcriptomics. BMC Genomics. 2018;19:477.

27. Stuart T, Butler A, Hoffman P, Hafemeister C, Papalexi E, Mauck WM 3rd, et al. Comprehensive integration of single-cell data. Cell. 2019;177:1888–1902.e21.

28. Golfinos-Owens AE, Lozar T, Khatri P, Johns ED, Hu R, Harari PM, et al. Integrated single-cell and spatial analysis reveals context-dependent myeloid-T cell interactions in response to immune checkpoint blockade in head and neck cancer. Clin Cancer Res [Internet]. 2026; Available from: 10.1158/1078-0432.CCR-25-2300

29. Andzinski L, Kasnitz N, Stahnke S, Wu C-F, Gereke M, von Köckritz-Blickwede M, et al. Type I IFNs induce anti-tumor polarization of tumor associated neutrophils in mice and human. Int J Cancer. 2016;138:1982–93.

30. Lopez-Leal F, Cabellos-Avelar T, Correa-Becerril DA, Juarez-Macias B, Cervantes-Diaz R, Reyes-Huerta RF, et al. Blockade of the CCR3 receptor reduces neutrophil recruitment to the lung during acute inflammation. J Leukoc Biol. 2024;116:1198–207.

