## Supplemental table and figures for "Single-cell multi-omics analysis reveals heterogeneity and plasticity of neutrophil states in response to immunotherapies"

**Supplementary Figure 1. Intratumoral injection of NAT components containing TNFα induces the formation of necrotic regions aross the tumor surface, which subsequently off, leading to complete tumor clearance in many mice.** Representative serial photographs of large M2h tumors taken at indicated time points following treatment in individual mice from: the untreated control group (Mouse F1, tumor-bearing), the aCD40 group (Mouse A1 and Mouse D3, both tumor-bearing), the aCD40 + TNFα group (Mouse G1, tumor-bearing and Mouse E2, tumor-free), and the NAT group (Mouse B1, tumor-free) treatment groups. Tumors are shown from the day of the first treatment dose (day 0) through the final imaging time point. Tumor-bearing mice show progressive tumor growth, with TNFα-containing treatment groups displaying visible necrotic lesions on the tumor surface in the days immediately following injection that differ from areas of central necrosis apparent in untreated control tumors that have outgrown their vascular supply. Notably, the necrotic surface patches induced by TNFα-containing therapies eventually fall off regardless of ultimate tumor outcome, although this detachment leads to complete and lasting tumor clearance only in a subset of mice. Tumor-free mice show gradual resolution of necrotic tissue and complete clearance of the tumor site by the end of the imaging period.

**Supplementary Figure 2. Treatment-Induced Tumor Clearance and Rechallenge Protection Following NAT-Based Therapy in M2h tumors.** (A and B) Average tumor volume (+ SEM) and overall survival of mice bearing small M2h tumors treated with ICI, NAT, or NAT + ICI. (C) Number of mice with sustained complete tumor clearance versus tumor-bearing mice at day 120 post implantation in each small tumor treatment group. (D and E) Average tumor volume (+ SEM) and overall survival of mice bearing large M2h tumors treated with aCD40 or NAT with or without. (F) Number of mice with sustained complete tumor clearance versus tumor-bearing mice at day 150 post implantation in each large tumor treatment group. (G and H) Overall survival of mice that cleared small or large M2h tumors after treatment and remained tumor-free at rechallenge, compared with age-matched naïve controls. TF = Tumor Free. For all statistical comparisons, $p<0.05$ = *; $p<0.01$ = **; $p<0.005$ = ***; $p<0.001$ = ****.

**Supplementary Figure 3. Label transfer of large M2h tumor neutrophil states onto small M2h tumor neutrophils shows similar overall trends of shifts in neutrophil state proportions across treatment conditions.** (A) Neutrophils were subsetted out *in silico* from a small M2h tumor scRNA-seq dataset, and subset identities (N0-N4) were assigned via label transfer using the large M2h tumor neutrophil annotations from Figure 2 as a reference. UMAP of small M2h tumor neutrophils colored by label transfer-predicted subset identity. (B) Stacked bar chart showing the proportions of each predicted neutrophil subset across treatment groups: control (untreated), aCD40, TNFα, Cetuximab, and NAT. (C) UMAPs as in (A), split by treatment group, providing a visual representation of the neutrophil subset compositional shifts quantified in (B).

**Supplementary Figure 4*.* Representative flow cytometry gating strategy for neutrophil state identification in large M2h tumors.** Flow cytometry gating plots used in FlowJo show the sequential gating strategy applied to large M2h tumor samples from separate *in vivo* experiments, progressing from scatter-based cell identification and doublet exclusion, through live cell and CD45+ selection, to the final identification of each neutrophil state based on the markers described in Figure 3.

**Supplementary Figure 5. KEGG pathway analyses predict distinct functional roles for N0-N4 neutrophils within large M2h tumors.** (A) Heatmap of normalized enrichment scores for selected KEGG metabolism pathways across neutrophil subsets, with red indicating pathway activation and blue indicating suppression. (B) Heatmap of normalized enrichment scores for selected KEGG apoptosis pathways across neutrophil subsets, with red indicating pathway activation and blue indicating suppression.

**Figure T1. Table of antibodies used for flow cytometry experiments.** Table listing all antibodies used for flow cytometric validation of neutrophil states in large M2h tumors, including target antigen, fluorophore, company, catalog number, clone, and volume used per test (µL). This panel was used across two independent *in vivo* experiments.

**Supplementary Figure 1.**

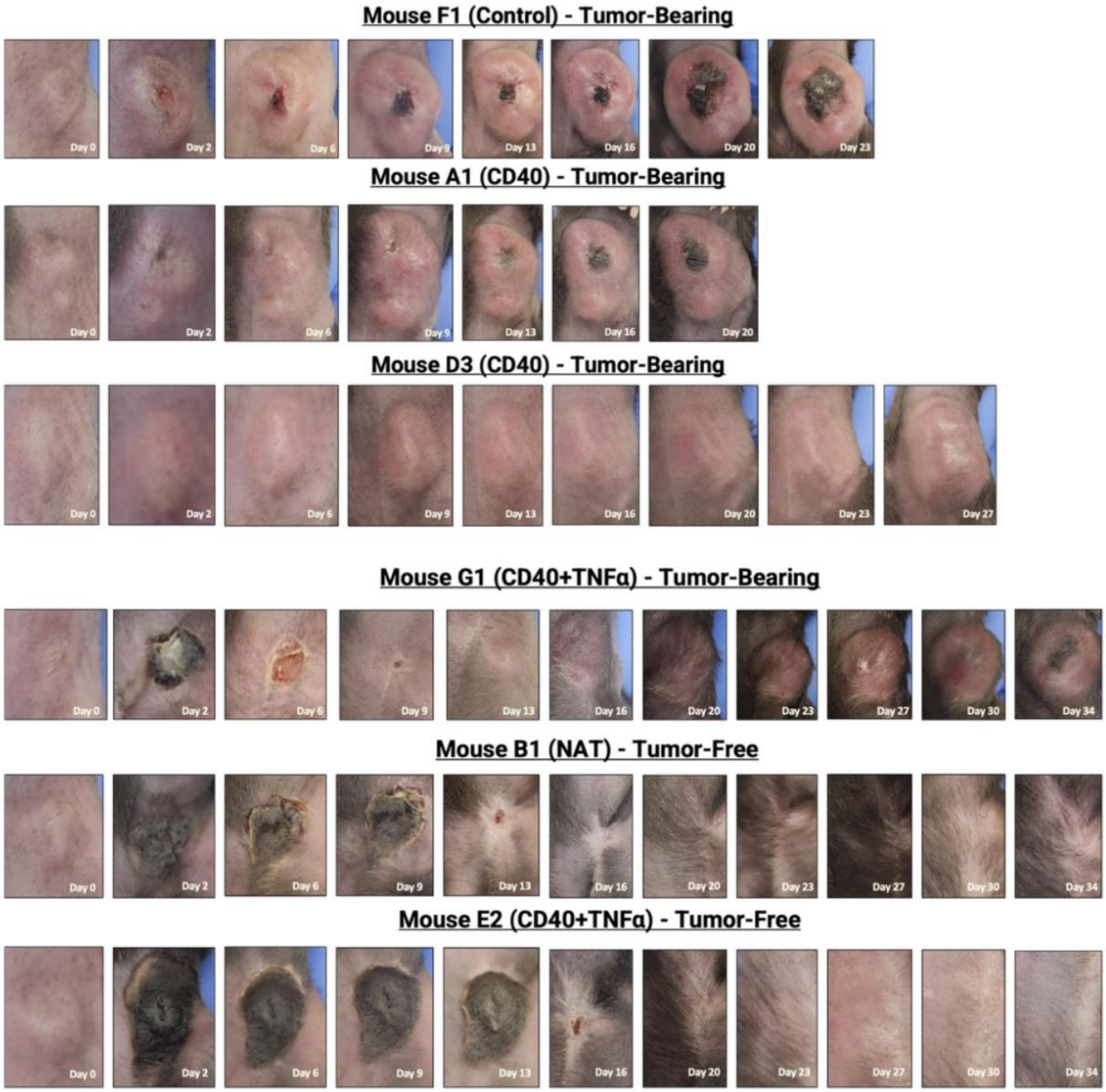

**Supplementary Figure 2.**

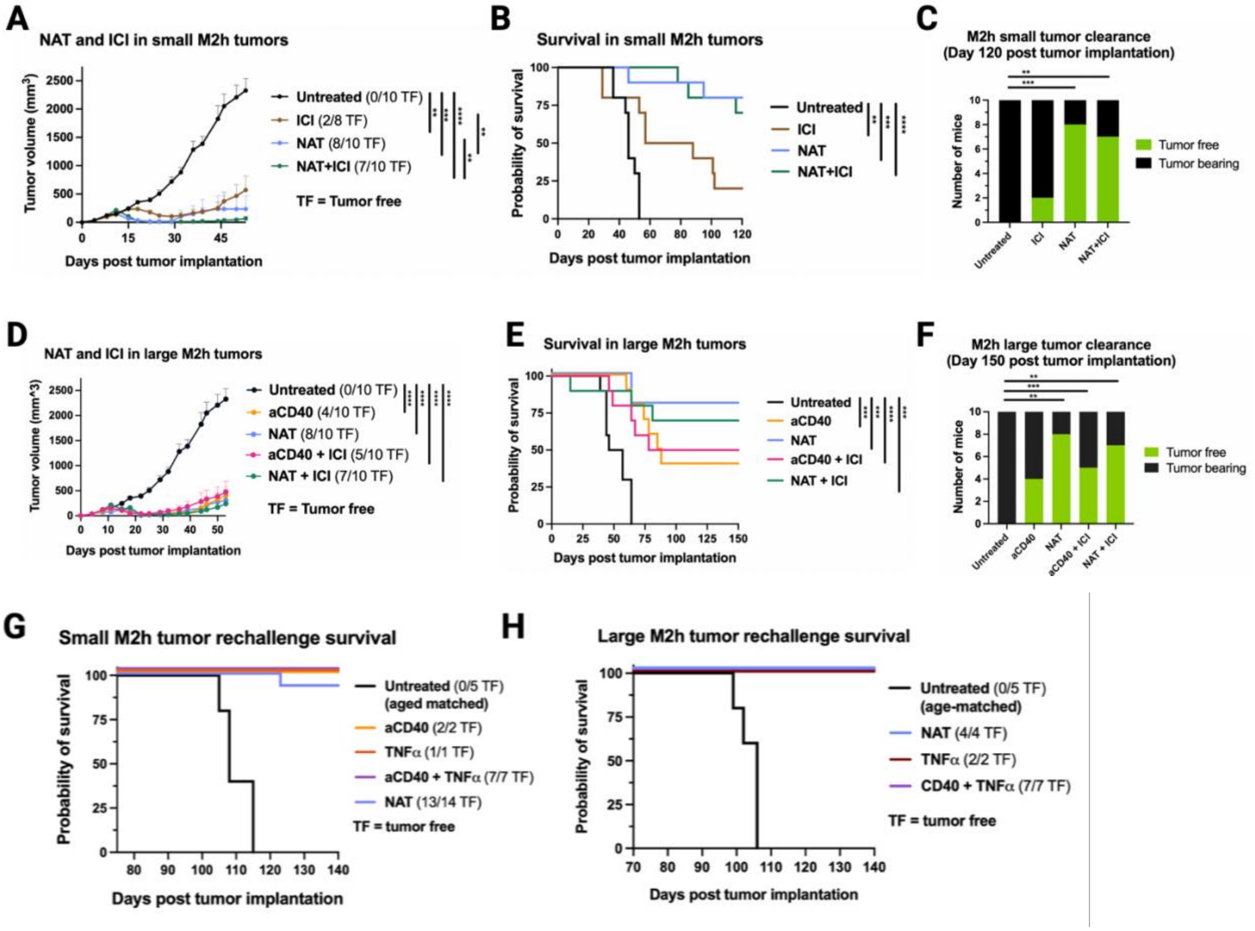

**Supplementary Figure 3.**

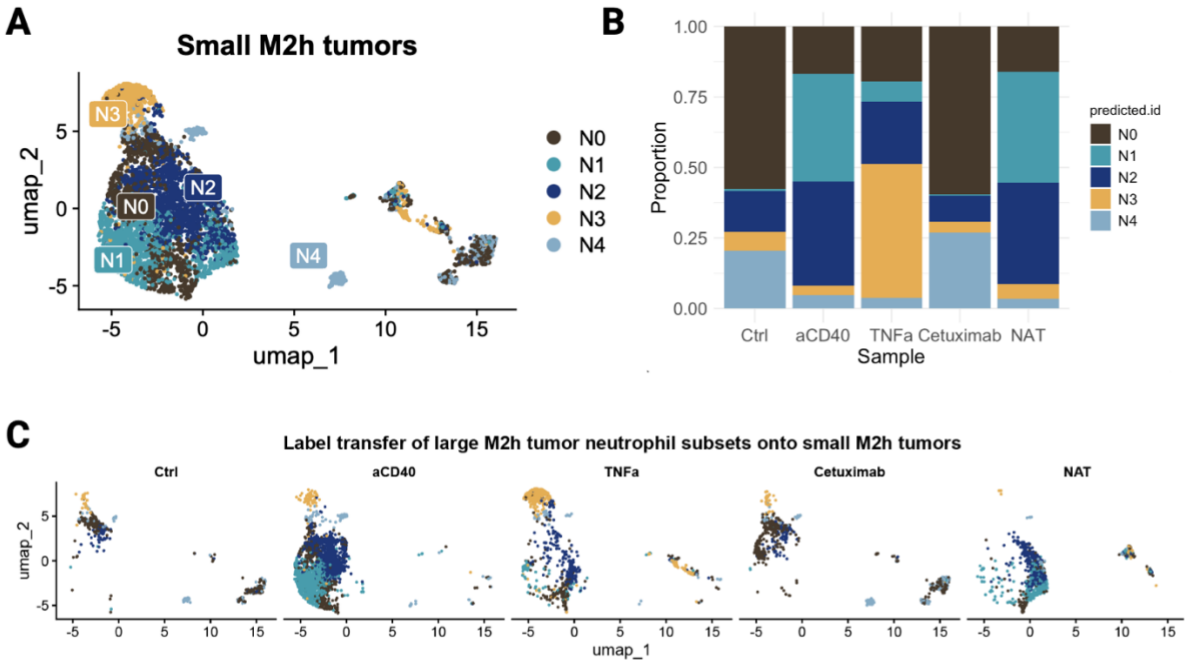

**Supplementary Figure 4.**

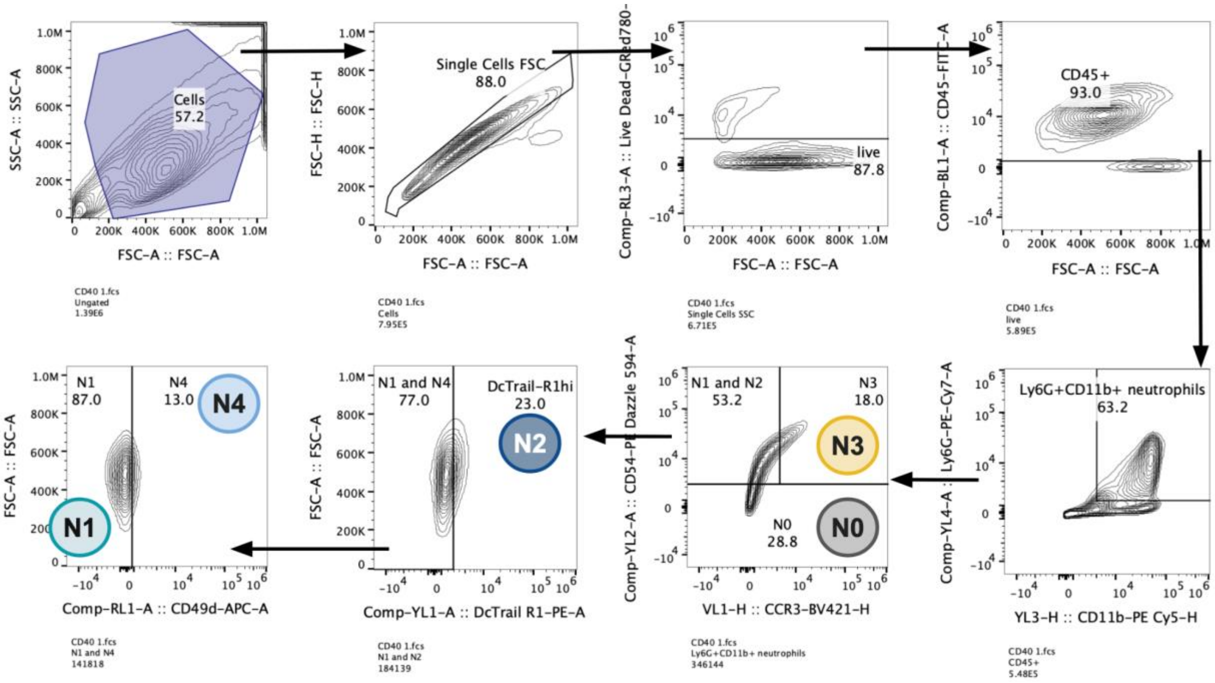

**Supplementary Figure 5.**

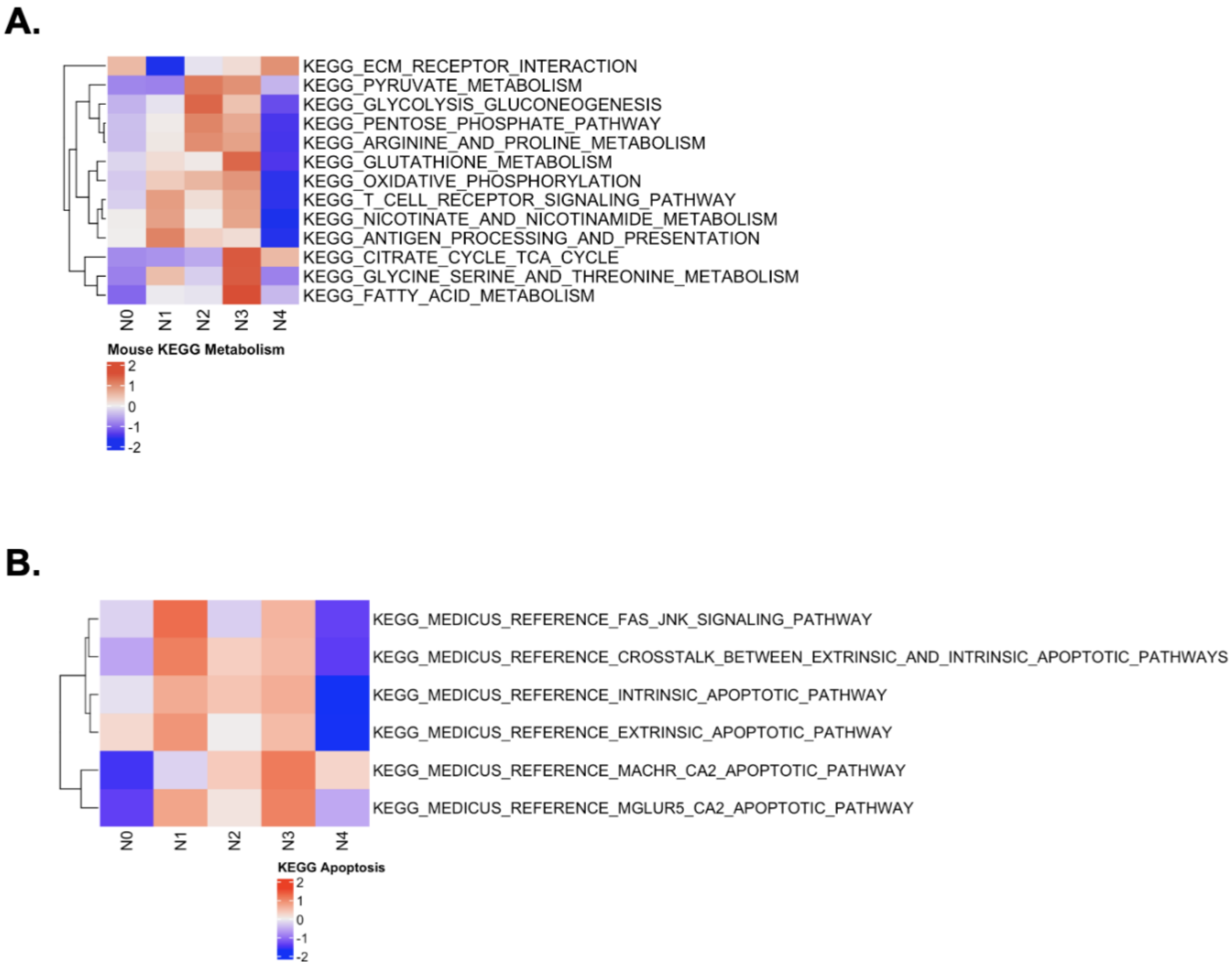

**Supplementary Table 1**

| Target | Fluorophore | Company | Catalog Number | Clone | Volume per test (μL) |
|---|---|---|---|---|---|
| CD45 | FITC | BioLegend | 103108 | 30-F11 | 1 |
| CD11b | PE-Cy5 | BioLegend | 101256 | M1/70 | 1 |
| Ly6G | PE-Cy7 | BioLegend | 127618 | 1A8 | 1 |
| Ly6C | BV605 | BioLegend | 128036 | HK1.4 | 2 |
| CD54 | PE-Dazzle 594 | BioLegend | 116130 | YN1/1.7.4 | 2 |
| CXCR2 | BV711 | BioLegend | 747812 | V48-2210 | 5 |
| CCR3 | BV421 | BioLegend | 144517 | J073E5 | 5 |
| DcTRAIL-R1 | PE | BioLegend | 133804 | mDcR1-3 | 5 |
| CD49d | APC | BioLegend | 103622 | R1-2 | 5 |
| MHCII | BV510 | BioLegend | 107636 | M5/114 | 5 |
| Viability | Ghost Dye Red 780 | Cytek | SKU-13-0865-T500 | N/A | 1 |
